# Ratiometric quorum sensing in *Enterococcus faecalis* conjugation

**DOI:** 10.1101/2019.12.11.871699

**Authors:** Alvaro Banderas, Arthur Carcano, Elisa Sina, Shuang Li, Ariel B. Lindner

## Abstract

Plasmid-mediated horizontal gene transfer of antibiotic-resistance and virulence in pathogenic bacteria underlies a major public health issue. Understanding how, in the absence of antibiotic-mediated selection, plasmid-bearing cells avoid being outnumbered by plasmid-free cells, is key to developing counter strategies. Here we quantified the induction of *Enterococcus faecalis*’ plasmidial sex-pheromone pathway to show that the integration of the stimulatory (mate-sensing) and inhibitory (self-sensing) signaling modules from the pCF10 conjugative plasmid, provides a precise measure of the recipient-to-donor ratio, agnostic to variations in population size. Such ratiometric control of conjugation favors vertical plasmid-transfer under low mating likelihood and allows activation of conjugation functions only under high mating likelihood. We further show that this strategy constitutes a cost-effective investment into mating effort, as overstimulation produces unproductive self-aggregation and reductions in the growth rate. A mathematical model suggests that ratiometric control of conjugation limits the spread of antibiotic resistance in absence of antibiotics, predicting a long-term stable co-existence of donors and recipients. Our results demonstrate how population-level parameters can control transfer of antibiotic-resistance in bacteria, opening the door for biotic control strategies. Ratiometric sensing in bacteria mirrors sexual behaviors observed in eukaryotes.

## Introduction

Horizontal gene transfer mediated by conjugative plasmids and integrative conjugative elements (ICEs) is a major cause for the rapid spread of bacterial antibiotic resistance (Johnson and Grossman, 2015). The process starts with a “mating” stage which depends on contact through sexual pili or cell-cell aggregation proteins followed via a type IV secretion system (Wallden, 2010). Importantly, expression of these conjugation determinants bears a significant fitness cost, reducing the host’s growth rate, implying the existence of a trade-off between the horizontal and vertical modes of plasmid transfer (Levin and Lenski, 1983). Indeed, across Gram-negative and Gram-positive bacteria, plasmid-donor populations are generally in a constitutive “off-state”, and only after induction by environmental factors or signaling molecules, is conjugation activated (Koraimann and Wagner, 2014). A general feature of these systems is that only a few members of the population activate the response (Koraimann and Wagner, 2014). This property suggests that antibiotic-resistant plasmids and ICEs naturally maintain a high proportion of vertical rather than horizontal transfer. On an evolutionary timescale, under a constant low plasmid-recipient availability, plasmid variants with lower conjugation rates are expected to gain a fitness advantage by increasing their vertical inheritance through host proliferation. Contrary to that, evolution is expected to favor plasmid variants with increased conjugation rates under conditions of constant high recipient availability (Turner et al, 1998). Therefore, the optimal effort that a plasmid invests on horizontal spread depends principally on the social environment under which conjugation control evolved. However, the primary population parameters determining the potential donor-recipient encounter rates, i.e. the densities of donors and recipients, can vary dynamically at faster timescales through growth, dilution and migration processes. Intuitively then, plasmid variants able to dynamically monitor mating likelihood and regulate conjugation effort accordingly, can evolve.

Antibiotic-resistance plasmids from *Enterococcus faecalis* are a major threat to public health (Klevens 2007) due to the efficiency of their pheromone-sensitive conjugation systems (Clewell, 2002) and the multiplicity of resistances they can transfer, including against the last-line antibiotic vancomycin (Murray 1997). Plasmids from *E. faecalis* exhibit the capacity to sense recipient densities (Bandyopadhyay et al, 2016). In the pCF10 plasmid and its family members, this mate-sensing is achieved with a unidirectional signaling (Dunny and Berntsson, 2016). This system is also no exception in terms of the cellular cost of conjugation, given the strong protein expression upregulation that ensues. Indeed, efficient cell-cell aggregation in this system relies on the high abundance of aggregation protein Asc10 (Christie et al,1988; Olmsted et al, 1993) as well as on the expression of >20 pheromone-regulated genes, including the ATP-dependent type IV conjugation machinery (Li et al 2012, Chen et al 2008). Donor populations exposed to a given level of inducer also have the capacity to sort into responding and non-responding cells, in this case through a pheromone-dependent bistable switch at the transcriptional level (Chatterjee et al 2011).

From an evolutionary perspective, sensing the presence of mates through selection for specificity (Fig1A, left and center) is straightforward to interpret as it avoids unproductive donor-donor interactions. Intriguingly however, in the *E. faecalis* system, two antagonistic plasmid-encoded pheromone-sensing systems control conjugation. These integrate information about the presence of potential plasmid recipients (mate-sensing) and about plasmid donors (self-sensing) (Fig. 1A, right; Chatterjee et al 2013). Such integration is antagonistic, with recipient-produced cCF10 and donor-produced iCF10 pheromones causing activation and repression of conjugation functions, respectively (Fig 1A, right). Such repressive self-sensing could effectively prevent self-aggregation and unproductive homophilic interactions at high donor densities, where donor-produced leaky cCF10 (Fig 1A) could accumulate to significant levels. Self-sensing is mediated by basal production of iCF10 during the growth of uninduced donors. We rationalized that self- and mate-sensing (Chatterjee et al 2013) could allow donor cells to perceive mate availability instead of mate numbers, impossible to achieve with classical quorum sensing (Mukherjee and Bassler, 2019), allowing cells to respond specifically to the population composition (Fig 1B). Intuitively, sexual aggregation or growth in liquid increases the concentration of both cCF10 and iCF10 (Fig 1B, right), but the effects produced by each could cancel each other out, producing a response that remains insensitive to fluctuations in the degree of cellular crowding. We demonstrate that such density-robust ratiometric control over horizontal plasmid transfer allows antibiotic-resistant *Enterococcus* plasmid-donors to estimate the conjugation likelihood in a cost-effective manner, and suggest that it robustly stabilizes the population composition in the long term, in the face of variation in resource availability.

**Figure 1.**
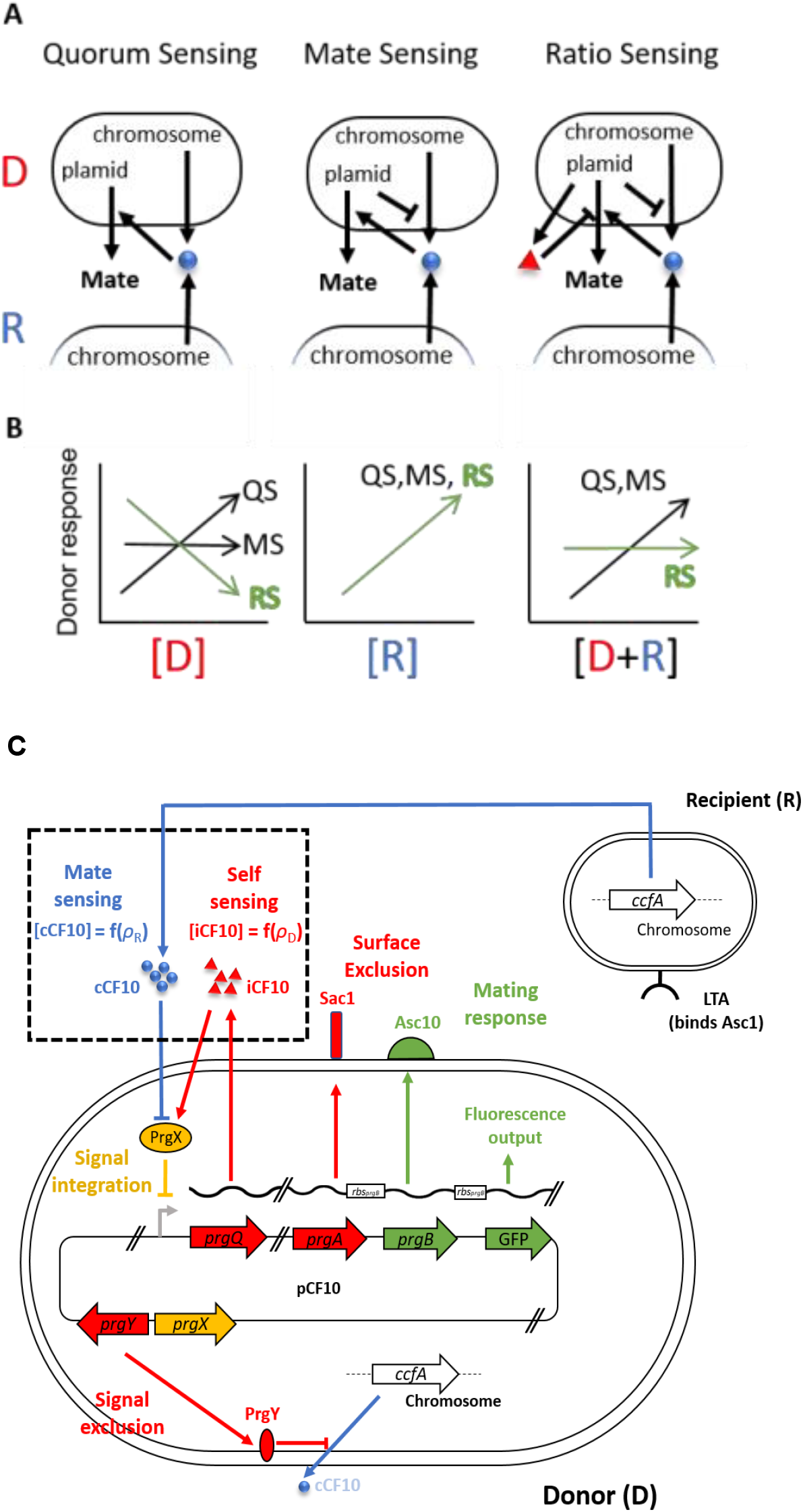
Population-parameter sensing in *Enterococcus faecalis* plasmid donors. **A.** Possible topological structures of pheromone-sensing in pCF10. Plasmids that perceive cCF10 pheromone (blue balls) from either donors or recipients (left) and consequently activate the pathway in proportion to the total population density (Quorum sensing) are less efficient at mating than variants that suppress the production of endogenous cCF10 (center, mate sensing). The present-day topological structure involves the presence of an antagonistic extracellular pheromone (iCF10, right). The extracellular concentration of cCF10 (blue balls) and iCF10 (red triangles) is proportional to the recipient and donor concentrations, respectively. **B.** The population-parameter disentanglement capabilities of pCF10 might explain the function of iCF10. Donor concentration ([D] (left), Recipient concentration [R] (center) and total population concentration [R+D] (right), are sensed differently by each one of the signaling schemes. Contrary to quorum sensing (QS) and mate-sensing (MS), only the ratio sensing (RS, green) scheme can distinguish specifically meaningful changes in recipient availability from simple fluctuations in crowding. **C.** The mating system of *Enterococcus faecalis*. The pCF10 plasmid in donor cells encodes “mating” (pre-conjugative) functions involved in self incompatibility (red), which allows avoidance of non-productive donor-donor interactions in at least three ways. First, by the Sec10 (encoded by *prgA*) activity which minimizes interactions of the neighboring, also cell-wall associated aggregation substance (Asc10, coded by the *prgB* gene) with lipoteichoic acid (LTA) in the cell walls of other donors (not shown) by steric hindrance (Dunny et al 1985), and keep those interactions specific for the LTA in recipients (shown). Second, by *prgY*, which minimizes production of the cCF10 pheromone (blue, Chandler et al 2005), a secreted product of the normal processing of a protein encoded by the *ccfA* gene, encoded in the genome which serves as the main cue used for activation. Finally, by secreting the iCF10 pheromone, which antagonizes the effect of cCF10 at the signal integration level, through competitive binding to the PrgX transcription factor (yellow). In this study, the pathway’s response was quantified by measuring Asc10-dependent phenotypes such as adherence to surfaces and sexual aggregate formation, and by monitoring the expression of a GFP reporter which is driven by a copy of *prgB*’s ribosome binding site (RBS) further downstream in the transcript (Cook et al 2011). Functions further downstream of *prgB* (including the conjugation machinery) are not shown.

## Results

### Ratiometric sensing of population composition

To test whether cells are indeed capable of distinguishing population-ratio from recipient density (Figure 1B), we quantified pheromone-mediated induction of the pCF10 plasmid during sexual aggregation by performing donor/recipient co-incubation experiments and measuring the gene expression response of the pCF10 mating-pheromone pathway (Fig 1C, see Methods) in plasmid donors. By homogenizing aggregated cultures (see Methods), we then used OG1RF(pCF10-GFP) donors (Cook et al, 2011), a fluorescent reporter strain of pheromone induction to both distinguish fluorescent donors from autofluorescent recipients and to quantify Asc10 expression in flow cytometry experiments (Fig. S1, see methods). For this, we resolved the dependency of the response on population-parameters (composition and total density) across a wide range of values (Fig 2A,B). We observed that ratio but not density causes activation of conjugation. This analysis showed donor cells commit to strong Asc10 expression only when populations are recipient-biased, and conversely suppressing the response in donor-biased populations. Crucially, the fraction of donor cells that show detectable response levels (Fig 2C) correlates with the result of mating assays (Fig 2D) in the wild type strain where on the one hand the mating efficiency reaches 100% (all donors mate once) at highly recipient-biased ratios and, on the other, remaining constant within a 5-fold difference in the total population cell-density after two hours, but remaining sensitive within two orders of magnitude of population-composition values. Our results show that the fraction of donors that mate increases at recipient-biased ratios but remains roughly constant when the total density is varied. This result implies that when populations are donor-biased, the activation of the system occurs in only a subset of cells during aggregation, consistent with the bistable model of pheromone induction (Chatterjee et al 2011) and further shows that donor population commits to conjugation only when in minority, and not at any specific recipient density, i.e. partner-sensing is ratiometric and not densitometric.

**Figure 2.**
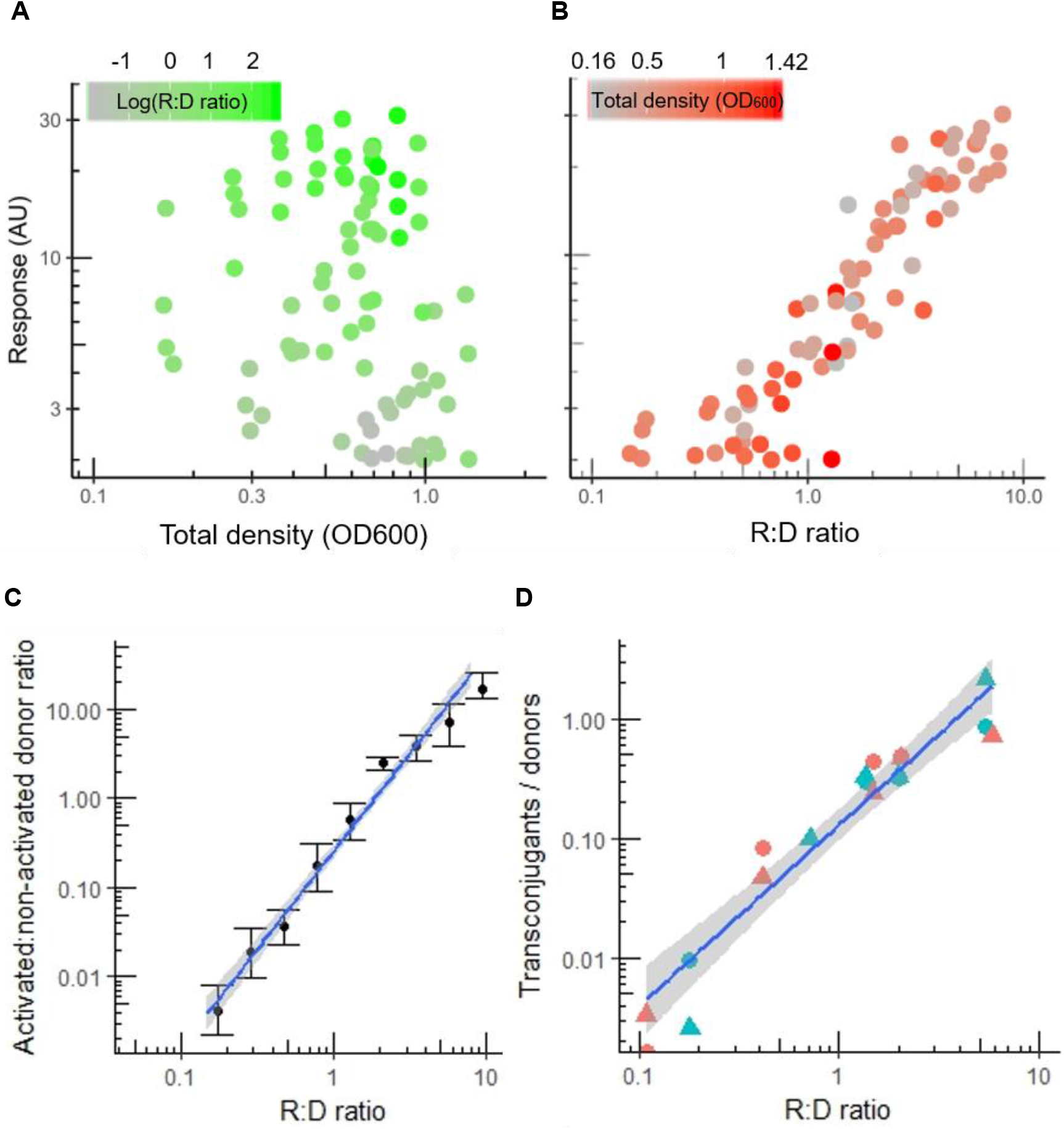
Population-density robust ratiometric sensing in *Enterococcus faecalis*. **A, B.** Response (mean GFP fluorescence intensity per cell) of plasmid pCF10 in populations with varying concentrations of emitter and donor cells at 2h after mixing and measured by flow cytometry (see methods). The response of OG1RF(pCF10-GFP) donors is plotted as a function of the two essential population parameters, the total population density [R+D] (A), and recipient to donor ratio [R:D] (B), each colored according to their counterpart parameter. **C-D.** The fraction of early (1h) activated donors as a function of the R:D ratio. Error bars are the SEM of 7 independent experiments. (C). The mating efficiency (see methods) as a function of the R:D ratio at 2 (circles) and 3 (triangles) hours with total densities (OD600) of 0.1 (red) and 0.5 (blue).

### Asc10 expression reduces vertical transfer of conjugative plasmids

Ratiometric sensing is reminiscent of the mating system of *S. cerevisiae*, where the population sex-ratio allows cells to sense the degree of mate-competition in the population (Banderas et al 2016). In yeast, A-type cells use sex-ratio sensing to balance a trade-off existing between commitment to mating and clonal haploid proliferation. However, whether ratiometric sensing relates to the existence of a similar trade-off in *E. faecalis* is unclear. To test whether a trade-off between investment in conjugation and vertical plasmid proliferation exists in *E. faecalis*, we first estimated the pheromone concentrations that correspond to the stimulation range observed in the co-incubation experiments. Shaken cultures of donors exposed to sufficient cCF10 concentration tend to self-aggregate, i.e. [cCF10]>10nM (Fig. S2). Although a reduction in growth below 10 nM is detectable by optical density measurements (Fig S3), growth is difficult to measure precisely under shaking conditions, either with optical density or CFU counts, as microaggregates are difficult to disperse. For this reason, we developed an assay to quantify growth in static cultures, where self-aggregation is not aided by orbital shaking and the pheromone-response is detected as Asc10-dependent surface attachment at concentrations below pathway saturation (Fig. 3A, see Methods). By quantifying growth under such conditions, we show that significant growth impairment occurs at pheromone concentrations marking the transition point to surface adhesion and that such impairment increases further at high pheromone concentrations (Fig. 3B, see methods). At concentrations in the μM range, we observed a time-dependent decrease in the optical density at the highest cCF10 concentrations, consistent with pheromone-induced toxicity (Bhatty et al, 2017). Importantly, at concentrations lower than ~1 pM, cells accumulate as an easily visible dense precipitate that can be easily resuspended, unlike Asc10-expressing cells which distributed homogeneously as a film in the bottom and remained adhered after resuspension (Fig. 3 A,B, see methods). This allowed us to determine that the physiological range of sensing, defined as the pheromone concentrations within which cells can activate Asc10 and avoid self-aggregation is roughly two orders of magnitude, consistent with the dynamic range of sensing on the recipient:donor ratio scale in coincubation experiments (Fig 2B).

**Figure 3.**
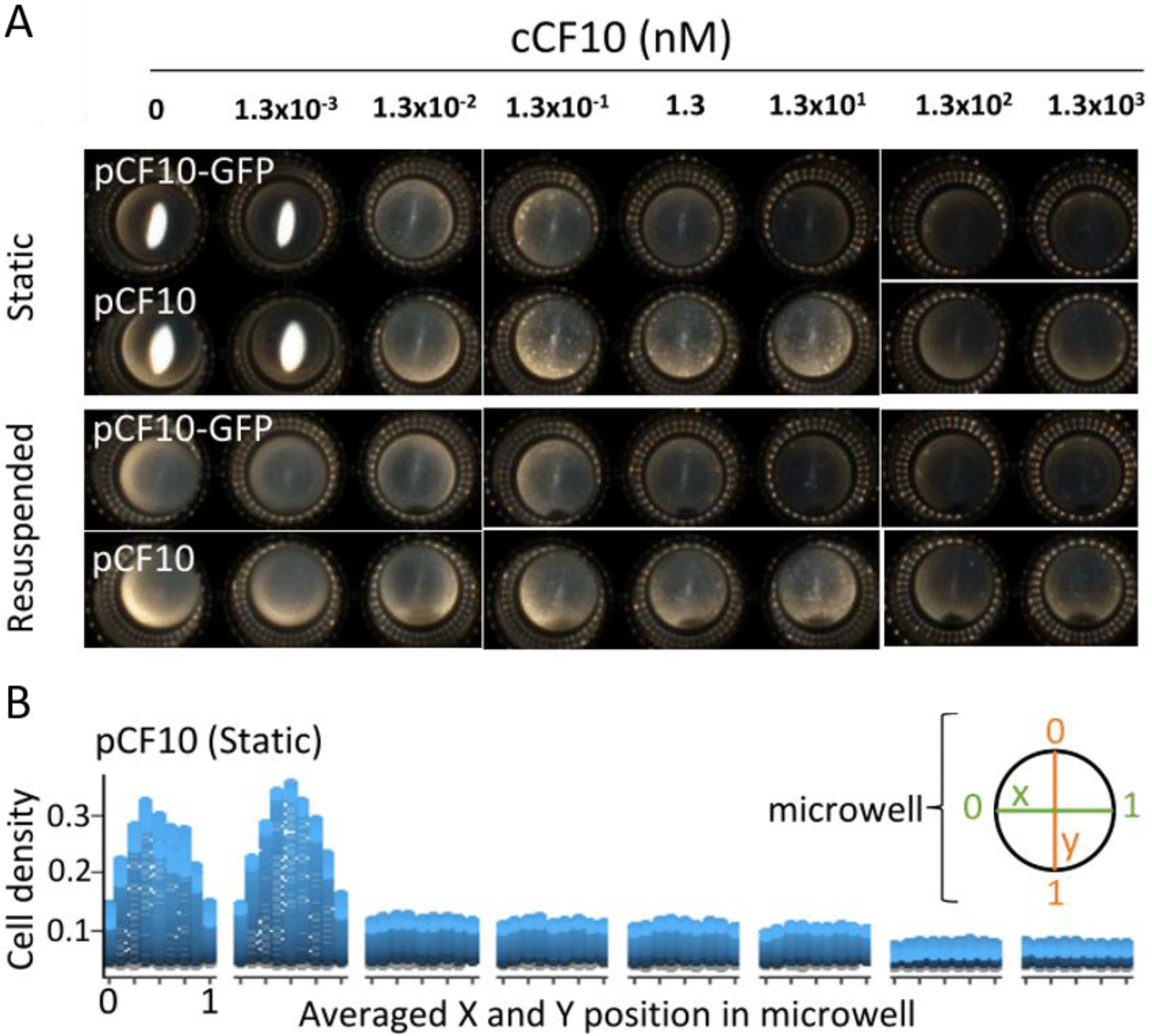
Response to purified cCF10 is sensitive and costly. **A.** A phenotypic assay for Asc10 activity (see methods) showing the macroscopic appearance of static cultures of OG1RF donors carrying wild type pCF10 or the pCF10-GFP Asc10 expression reporter (Cook et al 2011, see methods), after a 6-hour incubation period with varying concentrations of purified cCF10. Unstimulated cells appear as a precipitate in the center of the microwell. Adhered cells appear as a film. **B.** Time-resolved microwell spatial scanning of optical density in OG1RF(pCF10). Blue represents time from 0 (dark) to ~6 (light) hours.

Our results show that a trade-off between vertical and horizontal plasmid transfer exists in *E. faecalis*. This suggests that, similarly to the yeast case (Banderas et al 2016), specifically controlling Asc10 expression according to the recipient:donor ratio (a proxy for the mating likelihood) could be selected by evolution. Despite this, we also show that the degree of such pheromone-induced fitness loss has been partially reduced by the *prgU* gene (Bhatty et al 2017, Fig. S4, see methods), a putative RNA binding protein that negatively regulates pheromone-mediated induction and presumably ensues strong vertical transfer in the event of unsuccessful mating. Therefore, balancing the mentioned trade-off might not be the only advantage of ratiometric sensing. Indeed, any fitness-cost optimization strategy, is fundamentally limited by the inherent cost that donors pay for plasmid carriage, which will unavoidably favor recipient growth. For this reason, it has been suggested that natural *E. faecalis* populations could rather exist in a donor:recipient equilibrium (Chatterjee et al 2013). We therefore hypothesized that ratiometric sensing could play a role in the establishment and stability of such equilibrium.

### Mathematical model of conjugation dynamics

To understand the interplay between the discovered ratiometric regulation of conjugation and population stability, we built a simplified dynamic model consisting of donor and recipient populations described by two coupled ODEs. Donor population (*d)* is governed by three dynamics: first, an irreversible second-order mass-action-like process which depicts horizontal plasmid transfer (conjugation), transforming recipients (*r)* to donors with a rate constant (*λ*_*conj*_), weighed by function *h*(*r*, *d*) whose form depends on the strategy analyzed; second, a logistic growth law, with maximal rate *λ*_*d*_ limited by the carrying capacity *K* (*i.e.* vertical transfer); and third, an imposed constant dilution rate (*μ*):

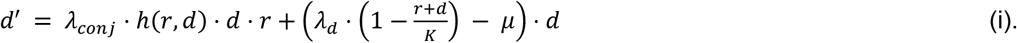

Likewise, recipients grow logistically with a maximal rate *λ*_*r*_, yet are removed from the pool by dilution and upon conjugation:

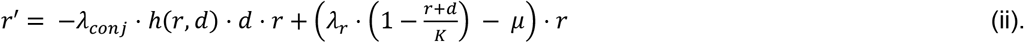

We assume that the presence of the plasmid burdens donor proliferation, *i.e.*, *λ*_*d*_ < *λ*_*R*_. Therefore, plasmid transfer decreases the fraction of recipients, and growth competition increases it. Using this approach, we compared four different strategies (*h*(*r,d*)): a strategy with constitutive activation (a constant value), an absolute (recipient only) sensing strategy (a sigmoidal function of the recipient density), an absolute (total population) sensing strategy (a sigmoidal function of the total density) and a relative (ratiometric) sensing strategy (a sigmoidal function of the population composition). Our results demonstrate that populations composed of donors and recipients can achieve a co-existence equilibrium point, independently of their initial composition (Fig 4A, FigS5). However, ratiometric sensing prevails as the only strategy allowing a robust co-existence of the two populations under wide fluctuations in resource abundance (*i.e.* carrying capacity *K*) (Fig 4B).

**Figure 4.**
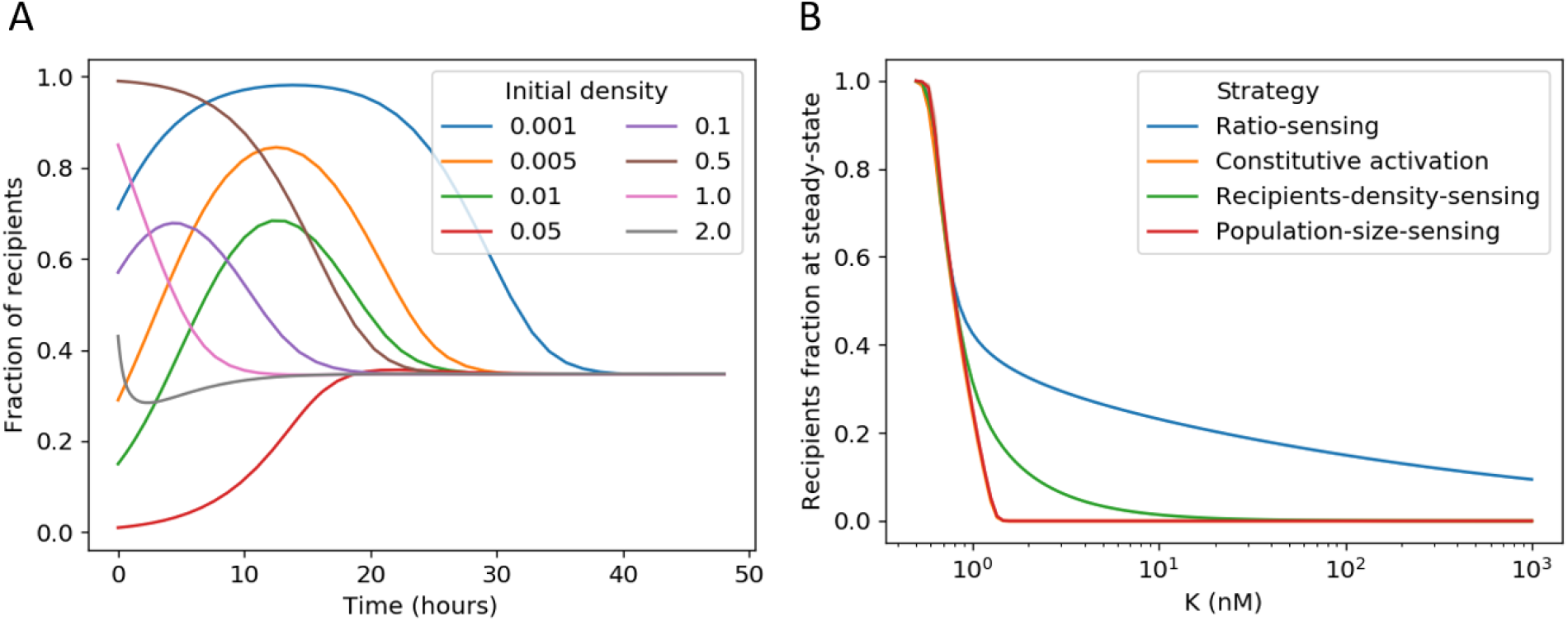
Mathematical model of mating population dynamics. **A.** Time-evolution of the system with different starting fraction of recipients and total population sizes. **B.** Steady-state recipient fraction for different sensing strategies as a function of the carrying capacity (*K*). Red and orange lines overlap.

Overall our modelling results provide a possible explanation for the observed absence of pure plasmid-bearing populations and suggest that non-trivial steady-state population dynamics can robustly arise in the absence of antibiotics and ratiometric population control.

## Discussion

Sexual aggregation and pheromone regulation are landmark properties of mating in single-celled eukaryotes but also operate in bacterial conjugation. Our results suggest that the latter further displays functional convergence with eukaryotic mating. The sex-pheromone pathways of *Saccharomyces cerevisiae* and *E. faecalis* share the same population-level property, sexual responses result from the antagonistic contributions of the cell’s own abundance (inhibitory) and that of its potential mates (stimulatory) (Chatterjee et al, 2013, Banderas et al, 2016), which produces phenotypic responses that depend specifically on the ratio between the two sub-populations.

Why does this ratiometric sensing evolve? The yeast example shows that it allows the “A” mating-type (one of the two mating types in *S. cerevisiae*) to balance a trade-off between its haploid growth (asexual reproduction) and mating effort (Banderas et al 2016). However, there is a fundamental difference between eukaryotic mating and bacterial plasmid conjugation: the latter is a spread and not fusion of genetic material. Then, as a selfish element, vertical spread of pCF10 is only limited by its cost to the host. Binary populations of antibiotic resistant and non-resistant bacteria with stable composition have been thought of as advantageous as a bet-hedging strategy in times of no antibiotic exposure (Chatterjee et al 2013). Our results point to the selective advantage of ratio sensing not only at the individual level (plasmid spread optimization by trade-off balancing), but also at the population levels: it predicts the establishment of a robust stable ratio, in line with experimental *in vivo* observations in a mouse model (Gary Dunny, personal communication). Such equilibrium is expected to be advantageous for the group under periodic antibiotic exposure. In their absence, growth is optimal and not limited by slower plasmid-bearing cells. However, recipients not taking over the population ensures survival of both plasmid and host when antibiotics are present.

An interesting connection emerges from evolutionary models where all mating systems (male/female in higher eukaryotes and mating-types in unicellular eukaryotes) share a common evolutionary origin, which is based on a system similar to bacterial conjugation (Hoekstra, 1990), namely, where an extra-chromosomal molecular symbiont (such as a conjugative plasmid) forces a population of asexual cells into a binary state (presence and absence of the plasmid, i.e. “sexes”) and mating (plasmid encodes the conjugation machinery for its spread, i.e. “sex”). Our theoretical results show that it is in principle possible to attain stability in such binary populations. Overall, our results show that ratiometric control over conjugation can limit the emergence of populations composed exclusively by resistant bacteria and provides clues on the early evolution of sexual systems.

## Acknowledgements

We thank Gary Dunny, Wei-Shou Hu, Ivan Matic, Dawn Manias, Rebecca Breue, Laura Cook, Christopher Kristich, Barbara Murray, Danielle Garsin, Lynn Hancock for valuable advice and/or strains. This work was financed by a LabEx *Who am I?* postdoctoral fellowship to Alvaro Banderas and the Fondation Bettencourt Schueller.

## Contributions

Conceived the project: AB; Designed research: AB, AL; Performed research: AB, AC, ES, SL; Wrote the paper: AB, AL. The authors declare no competing interests.

## Methods

### Bacterial Strains and Culture Conditions

For gene-expression quantification in co-incubation (mixed-population) experiments, the OG1RF(pCF10-GFP) reporter strain was used as donor (Cook et al 2011) and OG1RFSSp was used as the recipient cell. For mating experiments, strain OG1RF-GFP (expressing GFP constitutively, DebRoy et al 2012) was used as recipient and the wild type OG1RF-pCF10 was used as emitter. In general, overnight cultures were grown in Brain-Heart infusion media and appropriate antibiotics, whereas MM9YEG (Kristich et al 2007) semi-synthetic media was used for day cultures, inductions and co incubation experiments. The characterization of OG1RF(pCF10-GFP) as a useful reporter for coincubation experiments (See *coincubation experiments* below) was done by comparing it to a 2-fold dimmer wild-type GFP reporter strain OG1RF(pCF10-*prgC*-GFP) carrying a GFP downstream of *prgC* (a kind gift from Gary Dunny and Wei-Shou Hu).

### Pheromone stimulation

The cCF10 (synthesized by Biocat) pheromone was dissolved in 100% DMSO. Serial dilutions of cCF10 were prepared in DMSO and then added to 96 or 24-well plates (costar) containing the test cultures at an optical density of ~0.1 using an adjustable-spacer multichannel pipet or a multichannel pipet. Simulations have a constant final concentration of 0.1% DMSO (which does not affect growth, not shown) and 1% bovine seroalbumin (BSA, Sigma) to block peptide adhesion to surfaces. Before quantification, homogenization of self-aggregates was done mechanically by pipetting up and down ~50 times using a multichannel pipet, cultures where immediately diluted four times and sampling was performed quickly in a flow cytometer (see flow cytometry section)

### Coincubation experiments

After day culture growth, donor and receiver strains where mixed at different ratios and immediately serially diluted in 24-well plates, sealed (Breath-easy seals, Diversity Biotech) and shaken at 37°C (Infors orbital incubator). Strain OG1RF(pCF10-GFP) harbors a GFP construct located downstream of prgB (Cook et al 2011, Fig 1c) which is inserted within the prgU gene, as determined by sequencing (not shown). Since such mutation restricts growth (Fig. S4), by keeping GFP from dilution produced by cell division. This reporter shows an amplified output signal which is crucial to distinguish populations from autofluorescent recipients (Fig. S1). Also, the strain doesn’t mate efficiently further allowing us to minimize the effects of transconjugants that may alter the R:D ratio tested. The strain nevertheless maintains normal sexual aggregation and, importantly, remains sensitive in the same physiological range of pheromone stimulation as the wild type (Fig. S3). To calculate the fraction of stimulated donors we counted the stimulated donors (Fig S1, gate G1 plus gate G3) and divided it by the total amount of donors, which is the total amount of cells in the sample: [gate G1 + gate G2 + 2xG3 (each event in G3 is a cell pair)] multiplied by the donor fraction, which is a known variable *ab initio*.

### Flow cytometry

Flow cytometry was performed in a Gallios instrument (Beckman Coulter) or a Fortessa HT (Becton Dickinson) with sample sizes of 10000 to 1000000 cells. The fluorescence signal from GFP was acquired using a 488 nm laser for excitation, and a 525/40 (Gallios) or a 530/30 filter (Fortessa) for emission.

### Adherence/growth assays

Cell densities in the absence of aggregates were reconstructed from spatially resolved OD_600_ measurements done on glass-bottom micro-well plates (Corning) in a microplate reader (Tecan-Infinity, equipped with a monochromator) with an incubation temperature of 37°C, no shaking and the “lid on” mode activated. The lid of the plate was kept on and sealed with punctured parafilm to avoid evaporation. Twenty-eight positions within each microwell where acquired at regular intervals. After the experiment finalized, images were acquired with a SCAN 1200 colony counter. To avoid uneven illumination, three microwells per picture were acquired.

### Aggregation and turbidity assays

Aggregation levels were determined in 24-well plates, with shaking at 150 RPM at 37°C and BSA 1% in MM9YEG (Kristich et al 2007). Quantification was done by measuring the optical density decrease in time (Fig S3A). Aliquots were sampled near the liquid surface, touching the well wall with the pipet tip. The total turbidity loss (Fig. S3B) was done by sampling the culture at the last time point in Fig. S3A, after strong resuspension and an immediate dilution in MM9YEG.

### Mating experiments

Mating assays were performed by co-incubating the wild type OG1RF(pCF10) with constitutively fluorescent strain OG1RF-GFP (DebRoy et al 2012) for 2 or 3 hours under optimal conditions for aggregate formation, i.e. in 24-well plates (Costar) under orbital shaking at 150 RPM in a final volume of 1ml in the presence of 1% BSA. Homogenization of macroscopic sexual aggregates formed during mating reactions was done mechanically by pipetting up and down ~50 times using an adjustable-spacer multichannel pipet (Rainin) set to half of the total reaction volume (1 ml) and immediately diluting the samples to the appropriate dilution. Dilutions were plated in Brain Heart Infusion (BHI) agar medium. When colonies achieved an appropriate size, a velvet colony replicator was used to stamp the colonies on BHI containing tetracycline (10 μg/ml) to select for donors and transconjugants (and, by plate comparison, identify recipients), which we further distinguished from each other by detecting GFP positive colonies (transconjugants) in a transilluminator equipped with a camera (Biorad Chemidoc). The mating efficiency was calculated by dividing the number of transconjugants by the number of donors.

### Mathematical modelling

Coding was done on Python 3.7, with packages sympy 1.4, matplotlib 3.1.1, scipy 1.3.1. The “BDF” method of the ‘solve_ivp’ function was used to integrate ODEs. The different strategies were modeled as follows. First, for the ratiometric sensing strategy, we set 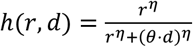. Mathematical analysis shows that, in this case, as long as *K*is larger than 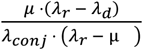, there exists an equilibrium point for this system, where both donors and recipients co-exist (Fig. 4). The carrying capacity (*K*) in our model with dilution is a proxy for environment quality. Hence, ratio sensing allows for the co-existence of both donors and recipients at the mere condition that the environment has enough quality. Second, for mating at a constant rate, if *h* equals zero, there is no mating, and because receivers grow faster than donors and there is dilution, they end up taking over the population. If *h* is a non-zero constant, we find that there is a non-trivial equilibrium where donors and recipients co-exist if and only if 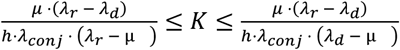. The lower bound is the same as in the ratiometric case but the existence of an upper bound makes the conditions more difficult to satisfy in practice, and the possibility of co-existence much less robust to environmental changes. Third, for the strategy with absolute-density sensing of mates, if 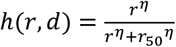 and knowing that the steady state population will be between 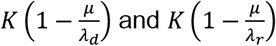, we can separate three cases. First, if 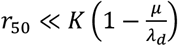 then *h* will be close to one for almost all possible ratios. Similarly, if 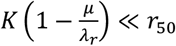, then *h* will be close to zero for almost all possible ratios. In these two cases, the system can be treated as in the previous paragraphs. If *r*_50_ is however of an order of magnitude between those of 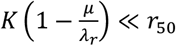 and 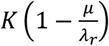, then the system behavior is not analysed here. In any case, absolute-density sensing of mates cannot be robust to variations in the quality of the environment, contrarily to ratio-based sensing. Finally, for the strategy where the total density of both subpopulations is sensed, we can have a similar reasoning as *h* will saturate for small and large K. Model (Fig. 4) parameter values are shown in Table 2.

**Table 1.** Strains used in this study

| *Enterococcus faecalis* strain | Description | Reference |
| --- | --- | --- |
| OG1RF(pCF10) | Wild type OG1RF carrying wild type pCF10 plasmid | Bourgogne et al, 2008 |
| OG1RF(pCF10-GFP) | pCF10 with a RBS<sub>*prgB*</sub>-GFP construct located between *prgB* and *prgC*. The cassette disrupts the small *prgU* open reading frame<sup>1</sup>. Alternative name: OG1RF(pCF10-LC1) | Cook et al, 2011 and sequencing results (not shown) |
| OG1SSp | Wild type recipient | Dunny et al, 1978 |
| OG1RF-GFP | Wild type expressing GFP constitutively. Alternative name is SD234 | DebRoy et al, 2012 |
| OG1RF(pCF10-*prgC*-GFP) | wild type pCF10 GFP construct located downstream of *prgC*. | Gift from Gary Dunny and Wei-Shou Hu |

<sup>1</sup>This ORF was unknown at the time of publication of the referred work.

**Table 2.** Model parameter values

|  |  |  |
| --- | --- | --- |
| $\lambda_{conj}$ | $0.012 \text{ nM}^{-1} \text{ min}^{-1}$ | From Chatterjee et al (2013) |
| $\theta$ | 0.5 | Arbitrary |
| $\eta$ | 4.3 | Fitted from figure 1 |
| $\lambda_D$ | $0.01548 \text{ min}^{-1}$ | From Chatterjee et al (2013) |
| $\lambda_R$ | $0.0201 \text{ min}^{-1}$ | From Chatterjee et al (2013) |
| $K$ | $2.0 \text{ nM}$ | Arbitrary |
| $\mu$ | $0.012 \text{ min}^{-1}$ | Arbitrary |

## Supplementary Material

**Fig S1.**
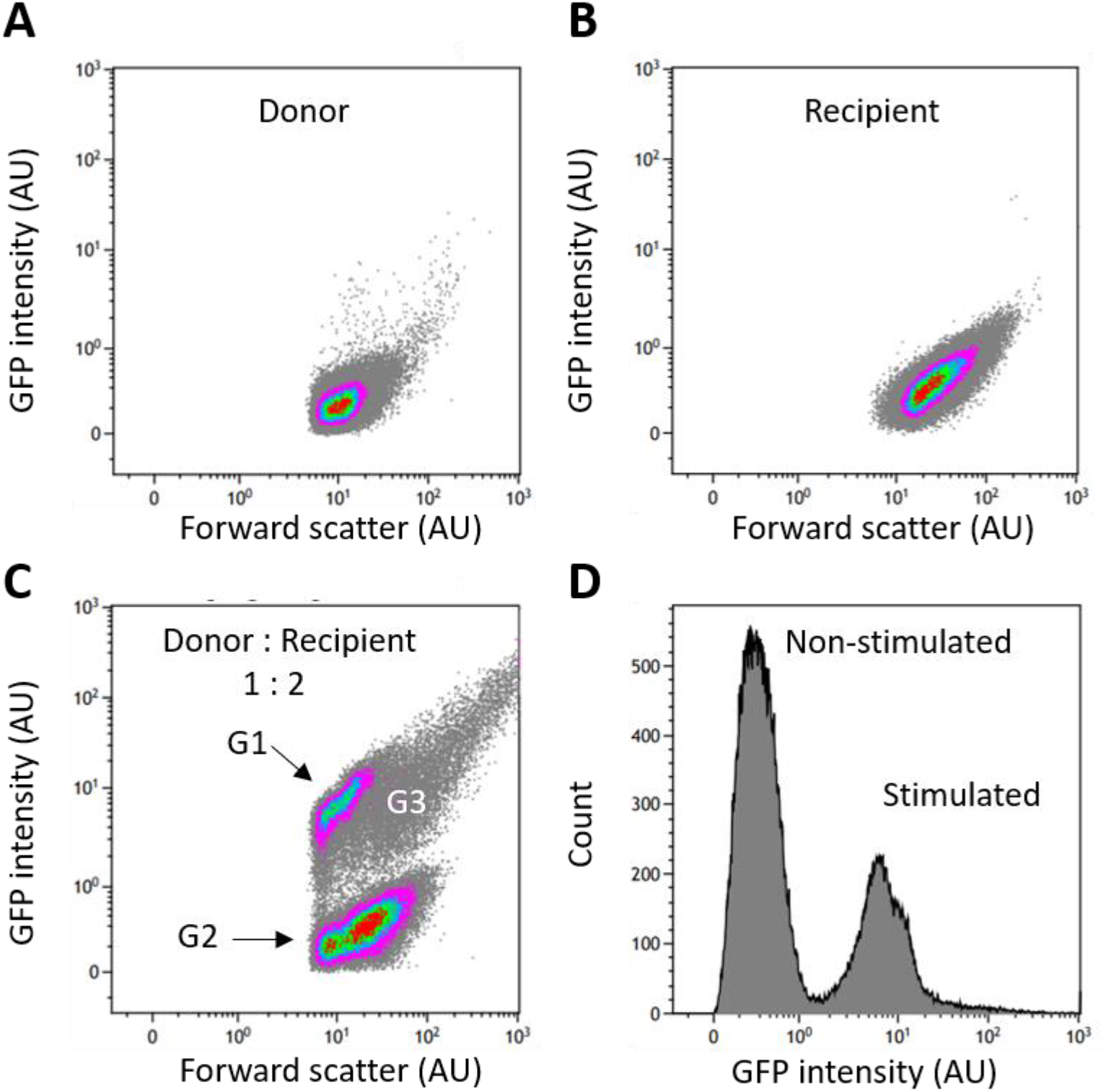
Identification of activated donors in mixed population experiments. A,B. Scatter plot of pure donor OG1RF(pCF10-GFP) (A) and recipient OG1SSp (B) cell populations. C. Example of a co-incubation reaction of donors and recipients at a recipient-biased ratio. The mean single-cell intensity of donors was obtained from population G1. Population G2 corresponds to a mixture autofluorescent donors and non-activated recipients. Population G3 corresponds to donor-recipient pairs (higher order aggregates, i.e. events in G3 with higher fluorescence values than events in G1 where not considered in the analysis). To estimate the fraction of activated donors (Fig. 2C) we considered the three populations and the known experimentally determined initial ratio in the calculation (see methods). D. Histogram with GFP intensities from the data on C.

**Fig S2.**
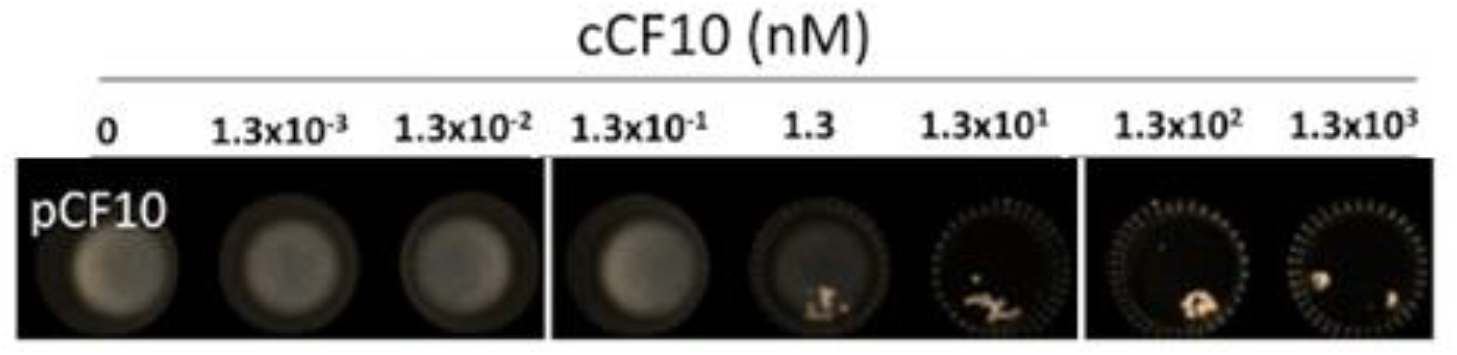
Orbital-shaking dependent macroscopic self-aggregate formation in OG1RF(pCF10) as a function of cCF10 concentration. The physiological (heterophilic) range of pheromone concentration required to induce detectable Asc10 expression lies below 1 nM cCF10.

**Figure S3.**
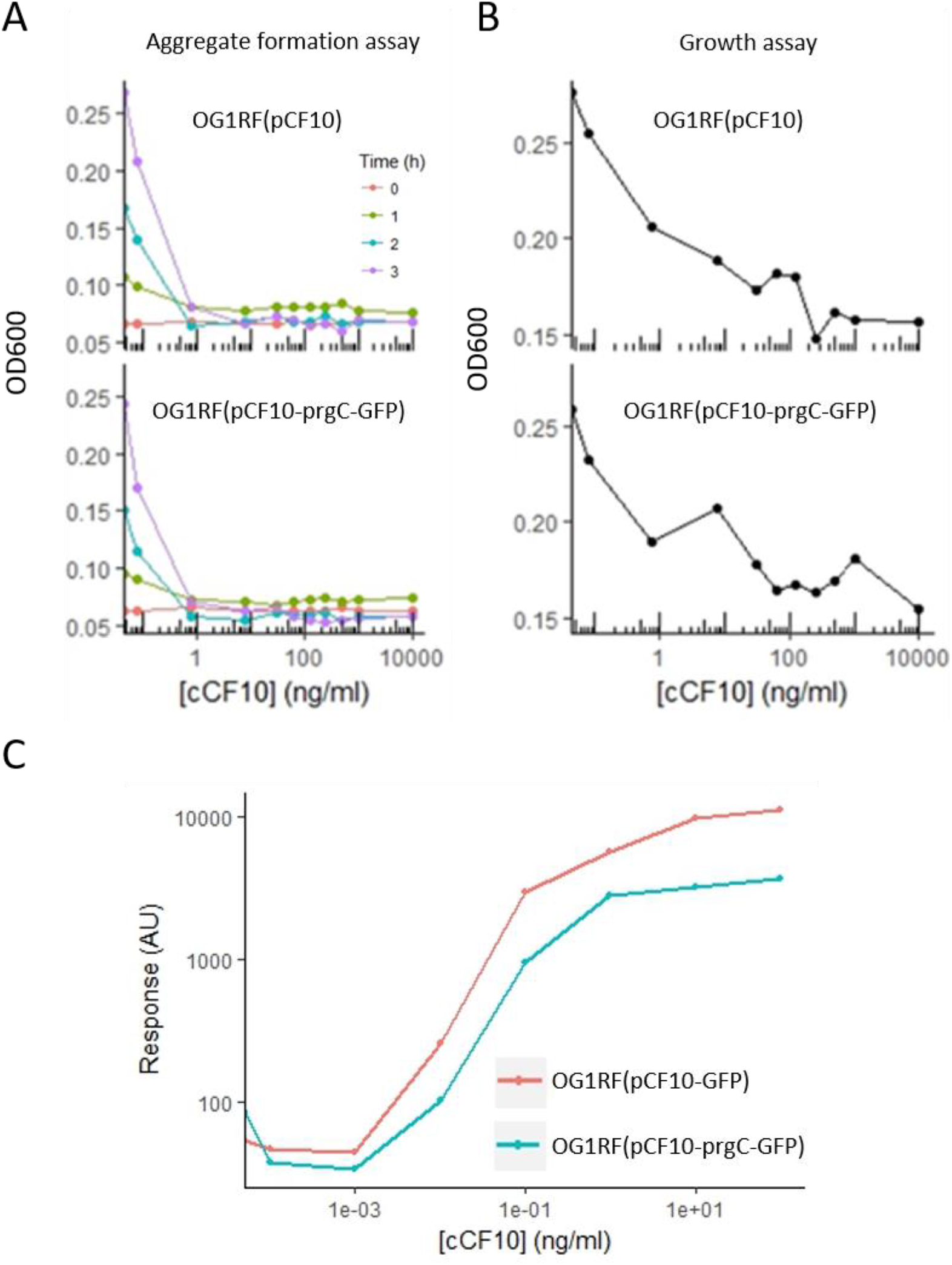
Gene expression reporter characterization. A,B. WT (OG1RF(pCF10)) versus OG1RF(pCF10-prgC-GFP) reporter comparison in an aggregate formation assay (A) and post aggregate-dispersion cell density measurement (B) (See methods). Note the increase in turbidity at 1h in A, time at which aggregation brings it back to the baseline. **C.** Comparison of Δ*prgU* GFP (OG1RF(pCF10-GFP)) and wild type (OG1RF(pCF10-*prgC*-GFP)) cCF10 dose-responses.

**Fig S4.**
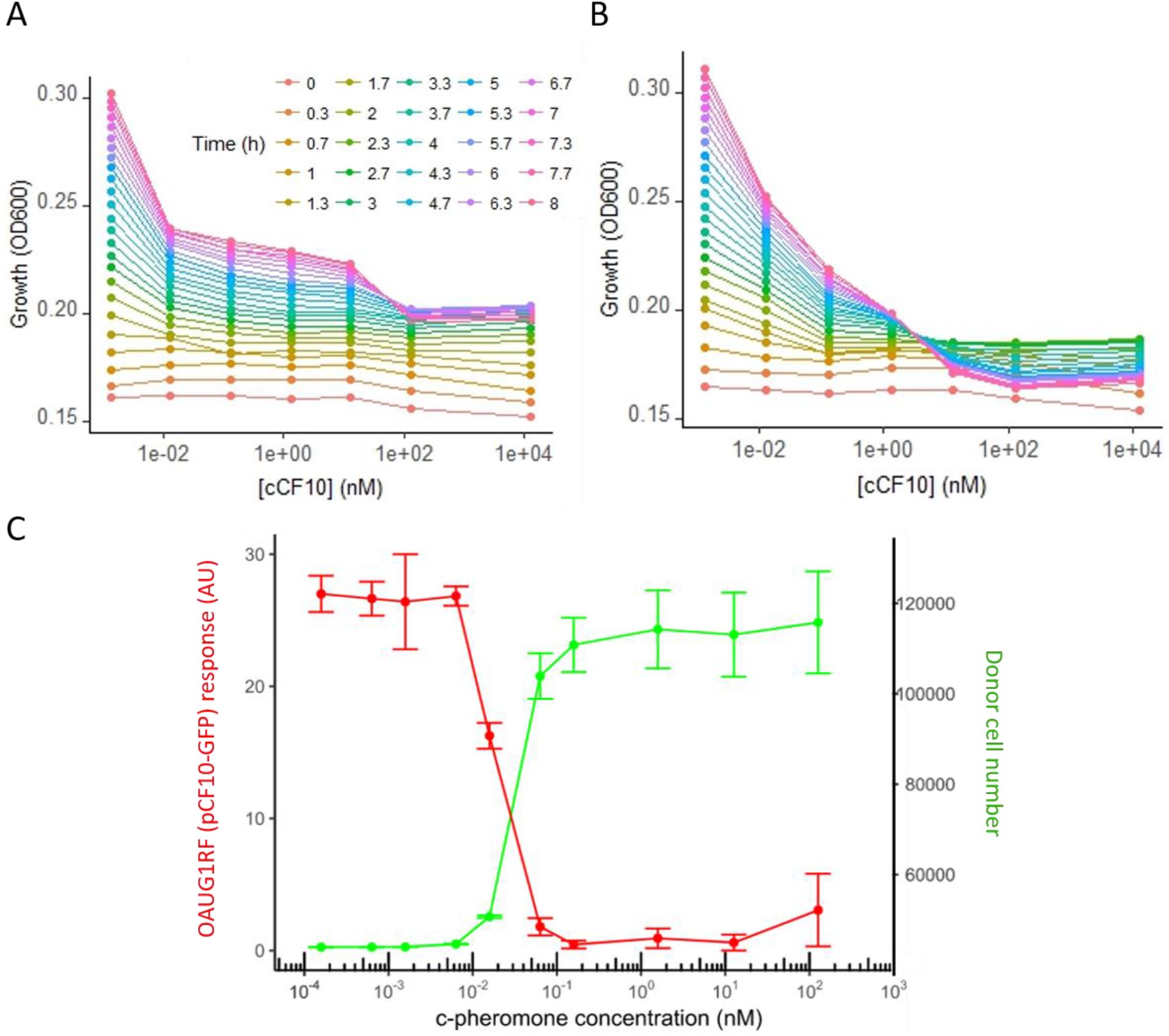
PrgU suppresses excessive fitness costs associated with unproductive activation of conjugation. A,B. Mean (per well) pCF10 (A) and pCF10-GFP (Δ*prgU*) (B) carrying donor growth in the phenotypic assay from (Fig. **3C**). **C.** Trade-off between growth and conjugation in the OG1RF(pCF10-GFP) reporter. Both growth and fluorescence intensity were obtained with a flow cytometer set to acquire constant sample volumes.

**Figure S5.**
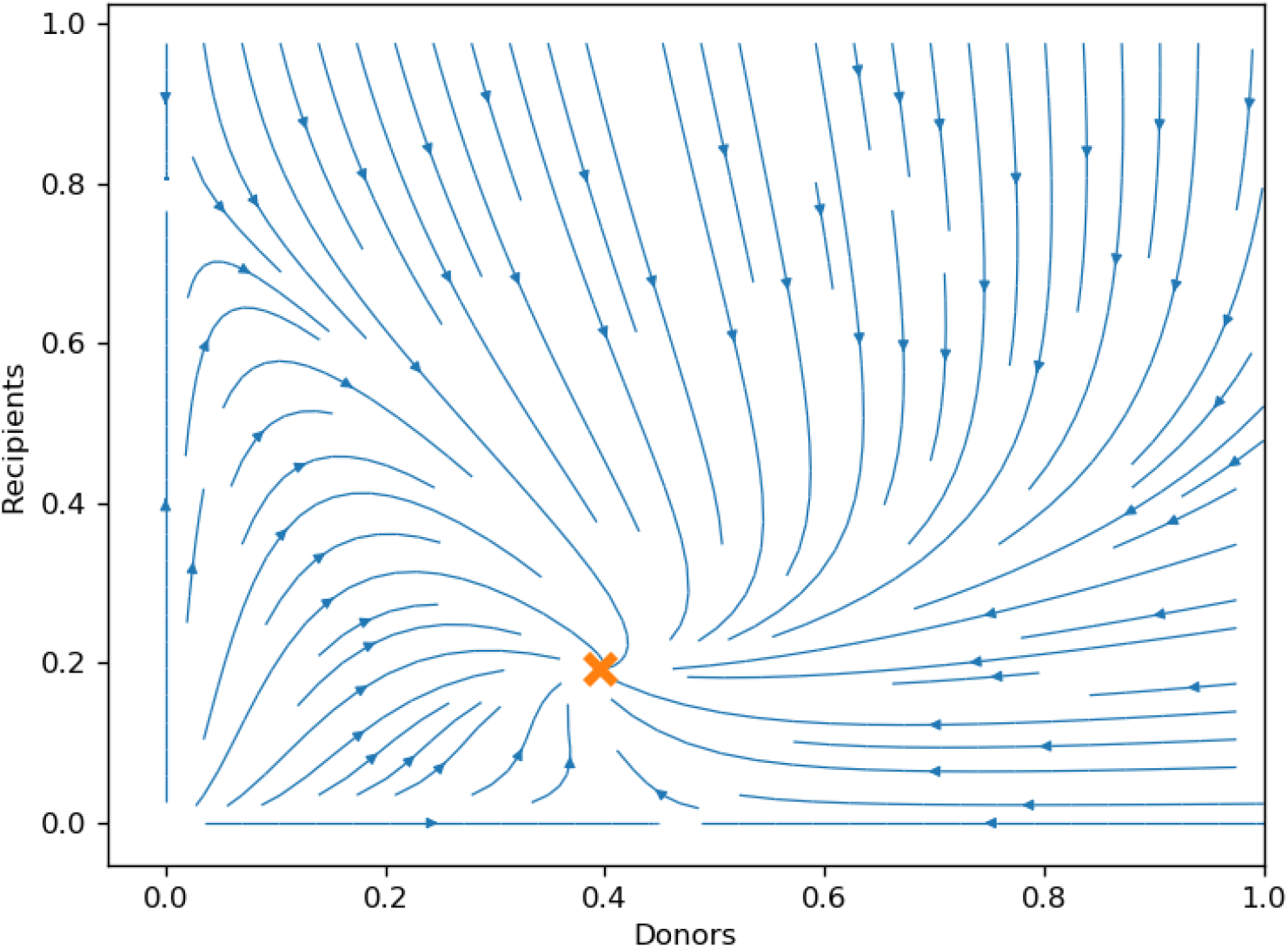
A stream-plot showing the time evolution (blue) of the simulated system dynamic converging towards a stable equilibrium recipient:donor composition (orange cross).

## Bibliography

1. Johnson, C. M., & Grossman, A. D. (2015). Integrative and conjugative elements (ICEs): what they do and how they work. Annual review of genetics, 49, 577–601.

2. Wallden, K., Rivera-Calzada, A., & Waksman, G. (2010). Microreview: Type IV secretion systems: versatility and diversity in function. Cellular microbiology, 12(9), 1203–1212.

3. Levin, B.R. & Lenski, R.E. (1983). Coevolution in bacteria and their viruses and plasmids Pp: 99–127 in Futuyma, D.J. & Slatkin M. Eds. Coevolution, Sinauer, Sunderland, MA.

4. Koraimann, G., & Wagner, M. A. (2014). Social behavior and decision making in bacterial conjugation. Frontiers in cellular and infection microbiology, 4, 54.

5. Turner, P. E., Cooper, V. S., & Lenski, R. E. (1998). Tradeoff between horizontal and vertical modes of transmission in bacterial plasmids. Evolution, 52(2), 315–329.

6. Klevens, R. M., Edwards, J. R., Richards Jr, C. L., Horan, T. C., Gaynes, R. P., Pollock, D. A., & Cardo, D. M. (2007). Estimating healthcare-associated infections and deaths in US hospitals, 2002. Public health reports, 122(2), 160–166.

7. Clewell, D. B., Francia, M. V., Flannagan, S. E., & An, F. Y. (2002). *Enterococcal* plasmid transfer: sex pheromones, transfer origins, relaxases, and the *Staphylococcus aureus*. Plasmid, 48(3), 193–201.

8. Murray, B. E. (1997). Vancomycin-resistant *Enterococci*. The American journal of medicine, 102(3), 284–293.

9. Bandyopadhyay, A., O’Brien, S., Frank, K. L., Dunny, G. M., & Hu, W. S. (2016). Antagonistic donor density effect conserved in multiple *enterococcal* conjugative plasmids. Applied and environmental microbiology, AEM-00363.

10. Dunny, G. M., & Berntsson, R. P. A. (2016). Enterococcal sex pheromones: evolutionary pathways to complex, two-signal systems. Journal of bacteriology, JB-00128.

11. Christie, P. J., Kao, S. M., Adsit, J. C., & Dunny, G. M. (1988). Cloning and expression of genes encoding pheromone-inducible antigens of *Enterococcus* (*Streptococcus*) faecalis. Journal of bacteriology, 170(11), 5161–5168.

12. Olmsted, S. B., Erlandsen, S. L., Dunny, G. M., & Wells, C. L. (1993). High-resolution visualization by field emission scanning electron microscopy of *Enterococcus faecalis* surface proteins encoded by the pheromone-inducible conjugative plasmid pCF10. Journal of bacteriology, 175(19), 6229–6237.

13. Li, F., Alvarez-Martinez, C., Chen, Y., Choi, K. J., Yeo, H. J., & Christie, P. J. (2012). *Enterococcus faecalis* PrgJ, a VirB4-like ATPase, mediates pCF10 conjugative transfer through substrate binding. Journal of bacteriology, 194(15), 4041–4051.

14. Chen, Y., Zhang, X., Manias, D., Yeo, H. J., Dunny, G. M., & Christie, P. J. (2008). Enterococcus faecalis PcfC, a spatially localized substrate receptor for type IV secretion of the pCF10 transfer intermediate. Journal of bacteriology, 190(10), 3632–3645.

15. Chatterjee, A., Johnson, C. M., Shu, C. C., Kaznessis, Y. N., Ramkrishna, D., Dunny, G. M., & Hu, W. S. (2011). Convergent transcription confers a bistable switch in *Enterococcus faecalis* conjugation. Proceedings of the National Academy of Sciences, 108(23), 9721–9726.

16. Chatterjee, A., Cook, L. C., Shu, C. C., Chen, Y., Manias, D. A., Ramkrishna, D., Dunny, G. M. & Hu, W. S. (2013). Antagonistic self-sensing and mate-sensing signaling controls antibiotic-resistance transfer. Proceedings of the National Academy of Sciences, 201212256.

17. Mukherjee, S., & Bassler, B. L. (2019). Bacterial quorum sensing in complex and dynamically changing environments. Nature Reviews Microbiology, 17(6), 371–382.

18. Cook, L., Chatterjee, A., Barnes, A., Yarwood, J., Hu, W. S., & Dunny, G. (2011). Biofilm growth alters regulation of conjugation by a bacterial pheromone. Molecular microbiology, 81(6), 1499–1510.

19. Banderas, A., Koltai, M., Anders, A., & Sourjik, V. (2016). Sensory input attenuation allows predictive sexual response in yeast. Nature communications, 7, 12590.

20. Bhatty, M., Camacho, M. I., Gonzalez-Rivera, C., Frank, K. L., Dale, J. L., Manias, D. A., … & Christie, P. J. (2017). PrgU: a suppressor of sex pheromone toxicity in *Enterococcus faecalis*. Molecular microbiology, 103(3), 398–412.

21. Kristich, C. J., Chandler, J. R., & Dunny, G. M. (2007). Development of a host-genotype-independent counterselectable marker and a high-frequency conjugative delivery system and their use in genetic analysis of *Enterococcus faecalis*. Plasmid, 57(2), 131–144.

22. Hoekstra, R. F. (1990). The evolution of male-female dimorphism: Older than sex? Journal of Genetics, 69(1), 11–15.

23. Chandler, J. R., Flynn, A. R., Bryan, E. M., & Dunny, G. M. (2005). Specific control of endogenous cCF10 pheromone by a conserved domain of the pCF10-encoded regulatory protein PrgY in Enterococcus faecalis. Journal of bacteriology, 187(14), 4830–4843.

24. Dunny, G. M., Zimmerman, D. L., & Tortorello, M. L. (1985). Induction of surface exclusion (entry exclusion) by Streptococcus faecalis sex pheromones: use of monoclonal antibodies to identify an inducible surface antigen involved in the exclusion process. Proceedings of the National Academy of Sciences, 82(24), 8582–8586.

25. DebRoy, S., van der Hoeven, R., Singh, K. V., Gao, P., Harvey, B. R., Murray, B. E., & Garsin, D. A. (2012). Development of a genomic site for gene integration and expression in Enterococcus faecalis. Journal of microbiological methods, 90(1), 1–8.

26. Dunny, G. M., Brown, B. L., & Clewell, D. B. (1978). Induced cell aggregation and mating in Streptococcus faecalis: evidence for a bacterial sex pheromone. Proceedings of the National Academy of Sciences, 75(7), 3479–3483.

27. Bourgogne, A., Garsin, D. A., Qin, X., Singh, K. V., Sillanpaa, J., Yerrapragada, S., … & Chen, G. (2008). Large scale variation in Enterococcus faecalis illustrated by the genome analysis of strain OG1RF. Genome biology, 9(7), R110.

